# Arbovirus persistence in mosquitoes is characterized by translation repression of viral RNAs

**DOI:** 10.64898/2026.02.26.708146

**Authors:** Marc Talló-Parra, Mireia Puig-Torrents, Gemma Pérez-Vilaró, Sol Ribó Pons, Juana Díez

## Abstract

Arboviruses induce acute lytic infection in human cells but establish persistent infection in their mosquito vectors, a viral strategy that is essential for sustained viral transmission. How mosquito cells maintain continuous production of viral progeny without compromising host cell viability remains a fundamental unresolved question. Because arbovirus replication in human cells relies on viral takeover of the host translational machinery, we investigated how translation is regulated during persistent infection in mosquito cells using chikungunya virus (CHIKV) as a model. A temporal analysis of viral RNA translation in RNAi-competent and RNAi-deficient *Aedes albopictus* cells revealed that persistence was associated with reduced viral protein production resulting from translation repression of viral RNAs. Subcellular localization analyses of the viral protein nsP2 and LC–MS/MS analyses of host tRNAs showed that, in contrast to human cells, CHIKV infection in mosquito cells neither induced nuclear relocalization of viral nsP2 to induce global host mRNA depletion, nor reshaped the tRNA modification landscape to compensate for the suboptimal codon usage of viral RNAs. Together, our results indicate that persistent infection in mosquito cells is characterized by a balanced host–virus translational state, in which limited viral translation is maintained while viral takeover of the host translational machinery is avoided. Notably, translation repression of viral RNAs was also observed during Zika virus (ZIKV) infection, suggesting that this mechanism may represent a general RNAi-independent feature of arbovirus persistence in mosquito cells.

## Introduction

Arboviruses transmitted by mosquitoes, including dengue (DENV), West Nile (WNV), Zika (ZIKV) and chikungunya (CHIKV) viruses, represent major global health threats [1]. In recent decades, urbanization, climate change, and globalization have facilitated the spread of these viruses and their mosquito vectors beyond tropical and subtropical regions, reaching new geographical areas worldwide. Consequently, more than 80% of the global population is currently at risk of infection by at least one of these viruses, with outbreaks increasingly reported in European countries [2].

Mosquito-borne viruses exhibit a remarkable ability to propagate efficiently in both human and mosquito hosts, two organisms separated by over one billion years of evolutionary divergence. In humans, these viruses cause acute and often severe infections, whereas in mosquitoes they establish persistent infections, a strategy essential for sustained transmission [3]. Although this persistence is typically compatible with long-term survival and does not cause overt pathology [4], reported effects on mosquito fitness including longevity, fecundity, and feeding behavior are heterogeneous and in some cases contradictory. These discrepancies likely reflect differences in mosquito and virus genotypes, infection route and dose, and environmental conditions [5–12].

How arboviruses replicate successfully in these evolutionarily distinct hosts while producing such distinct infection outcomes remains poorly understood. Moreover, molecular interactions between arboviruses and their mosquito hosts are far less well characterized than those described in vertebrates. Because persistent infection in mosquitoes helps ensure successful arbovirus transmission and propagation, understanding the mechanisms that support this state is critical for developing novel vector control strategies.

A key step in the arbovirus replication cycle is the efficient expression of viral proteins, which relies entirely on the host cell machinery [13]. Interestingly, the codon usage of diverse arboviruses, including CHIKV, reflects an intermediate usage pattern associated with replication in both vertebrate and mosquito hosts and is suboptimal when each host is considered in isolation. In both vertebrates and mosquitoes, highly expressed genes preferentially use G/C-ending codons, whose cognate tRNAs are highly abundant, whereas arbovirus genomes are enriched in A/U-ending codons, whose cognate tRNAs are comparatively scarce [14]. This codon bias is predicted to slow down translation elongation and decrease protein expression. However, using CHIKV as a model, recent studies show that in human cells, CHIKV infection overcomes this limitation by inducing alterations in the tRNA modification landscape. Specifically, CHIKV infection increases the levels of the 5-methoxycarbonylmethyluridine (mcm^5^) tRNA modification. This alteration reprograms codon optimality, selectively enhancing the translation of specific suboptimal codons that are highly enriched in the CHIKV genome. Hence, CHIKV codon usage becomes optimally matched to the tRNA modification landscape in infected human cells [15]. Moreover, in vertebrate cells, CHIKV non-structural protein 2 (nsP2) translocates to the nucleus, where it induces degradation of the RNA polymerase II subunit B1 (Rpb1), causing a massive shutdown of competitive host mRNAs [15,16].

Whether a similar strategy operates in mosquito cells, and how these cells sustain viral protein synthesis without compromising their own viability, remains unknown. Here, using CHIKV as a model, we address these questions by performing a temporal analysis of viral protein expression, translation dynamics, and tRNA modification landscapes. In contrast to human cells, we found that CHIKV persistence in mosquito cells is characterized by virus-specific and RNA interference (RNAi)-independent translational repression, enabling sustained yet limited viral protein production without major perturbation of host mRNA translation. Reduced CHIKV protein synthesis is associated with two main constraints: inefficient decoding of viral codons, as no changes in the tRNA modification landscapes were induced, and the subcellular behavior of the viral nsP2, which does not translocate to the nucleus in mosquito cells. We propose that a finely-tuned virus-host translational equilibrium supports both the sustained viability of mosquito cells and the establishment of arbovirus persistence through controlled viral protein synthesis.

## Results

### CHIKV persistence in mosquito cells is associated with reduced viral protein levels

CHIKV is a positive-sense single-stranded RNA ((+)ssRNA) virus whose genome structurally mimics host messenger RNAs (mRNAs), bearing a 5’ cap and a 3’ poly(A) tail. The CHIKV RNA genome (gRNA) contains two open reading frames (ORFs). The first ORF is translated directly from the genome and encodes the non-structural proteins (nsPs) required for viral replication, whereas the second ORF is expressed from a subgenomic RNA (sgRNA) transcribed during replication and encodes the structural proteins (sPs) necessary for virion assembly (**Fig 1A**). To investigate how CHIKV establishes persistent infection in mosquito cells without major effects on host cell physiology, we first analyzed CHIKV infection kinetics in *Aedes albopictus* U4.4 and C6/36 cells to identify the time point at which viral replication reaches a plateau, hereafter referred as persistence stage, defined by sustained viral RNA and infectious virus levels in the absence of cytopathic effects. U4.4 cells were selected as a physiologically relevant mosquito model with an intact antiviral RNAi response, whereas C6/36 cells, which lack a functional RNAi pathway, were included to determine whether the translational phenotypes observed during CHIKV persistence depend on a functional RNAi pathway. Both cell lines were infected with CHIKV LR2006-OPY1 strain at a multiplicity of infection (MOI) of 4, a condition under which the majority of cells become infected by 8 hours post infection (hpi) in C6/36 cells and by 1 day post infection (dpi) in U4.4 cells (**S1A Fig**). This strain was selected as it contains the *Aedes albopictus*-adapting mutation A226V. Samples were collected at early (3 and 8 hpi) and late time points (1, 2, 5 and 7 dpi). Viral titers (**Fig 1B**) as well as gRNA and sgRNA levels (**Fig 1C**) and their corresponding protein products (**Fig 1D-E**), were quantified over time. In parallel, cell proliferation and cellular ATP levels were monitored (**S1B-C Fig**).

**Fig 1.**
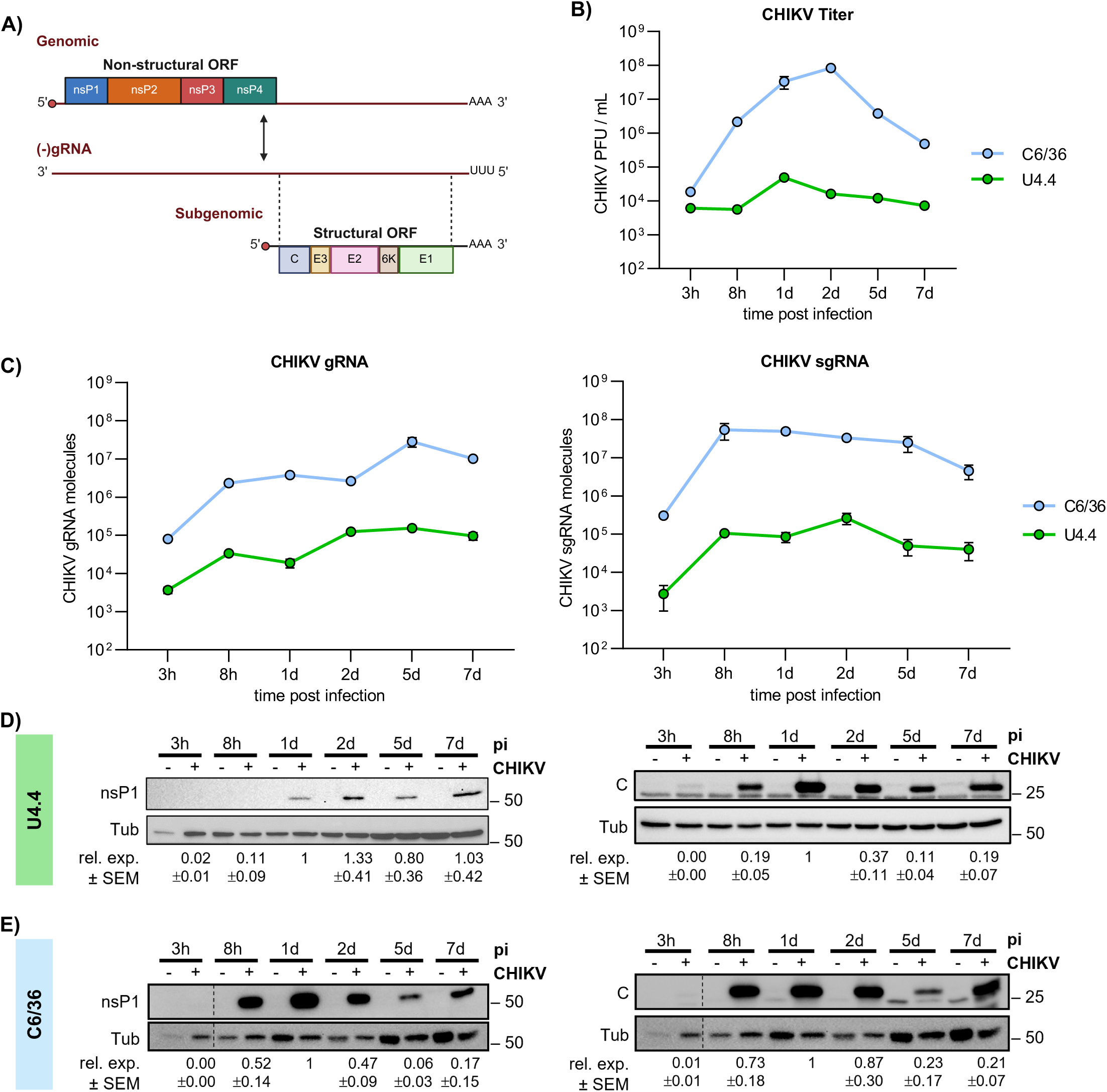
Kinetic of CHIKV RNA accumulation and protein expression in U4.4 and C6/36 mosquito cells. (A) Schematic representation of CHIKV RNA genome organization and viral protein expression strategy. (B) CHIKV titers over time in U4.4 and C6/36, determined by plaque assay. (C) Kinetics of CHIKV gRNA and sgRNA levels in U4.4 and C6/36, quantified by RT-qPCR. (D-E) Expression of CHIKV nsP1 and capsid (*C*) proteins in U4.4 cells (D) and C6/36 (E) over the course of infection. Green and blue colors correspond to U4.4 and C6/36 cells, respectively. All infections were performed at an MOI of 4 and samples were collected at the indicated time post infection. Data points represent the mean ± SEM of 3 independent replicates. Viral RNA levels were quantified using a standard curve generated from *in vitro*-transcribed CHIKV RNA. Protein expression levels correspond to band intensities normalized to tubulin and to the 1 dpi value within each blot. Dotted lines indicate separated lanes in the same membrane.

Comparative analyses of both cell lines showed that U4.4 exhibited lower viral titers, viral RNA levels and protein expression than C6/36 cells (**Fig 1B-E**), consistent with the presence of an active RNAi response. In U4.4, viral titers remained relatively stable throughout the time course, with a peak at 1 dpi. Similarly, but more pronounced, viral titers in C6/36 peaked at 1-2 dpi before declining. In parallel, intracellular viral RNA levels were determined by RT-qPCR. Both CHIKV gRNA and sgRNA levels increased over time and subsequently reached a plateau, after which only minor changes were detected. Specifically, positive-strand gRNA accumulated progressively in both cell lines and reached a plateau by 2-5 dpi in U4.4 and by 5-7 dpi C6/36 (**Fig 1C**). A similar pattern was observed for negative-stranded gRNA ((-)gRNA), except that in C6/36 the plateau was reached at 1 dpi (**S1D Fig**). In contrast, sgRNA plateaued rapidly at 8 hpi in both cell lines and subsequently remained relatively stable. Cell proliferation and cellular ATP levels were monitored throughout infection (**S1B-C Fig**). CHIKV infection induced an early arrest of cell proliferation and maintained ATP levels in C6/36 cells, but not in U4.4 cells. These changes correlated with the higher levels of viral replication observed in C6/36 cells and indicate that high-level CHIKV replication imposes a metabolic and proliferative burden on mosquito cells without causing cell death. We next assessed the expression of the viral proteins nsP1 and capsid (*C*) (**Fig 1D-E**). Notably, protein expression profiles did not mirror viral RNA kinetics. nsP1 expression in C6/36 and in U4.4 cells peaked at 1 dpi and 2 dpi, respectively, and then declined. Capsid expression peaked at 1 dpi in both cell lines and subsequently decreased. This discordance between declining protein levels and sustained viral RNA accumulation is consistent with post-transcriptional regulation, potentially involving inefficient translation of viral RNAs and/or their sequestration in compartments that are inaccessible to RNases and to the translational machinery.

### Translation of CHIKV RNAs is repressed in persistently infected mosquito cells

To directly determine whether the distinct CHIKV RNA and protein kinetics were linked to translation regulation, we performed polysome profiling analyses. This approach provides a high-resolution view of the translational landscape by resolving total mRNAs according to ribosomal occupancy, thereby distinguishing untranslated RNAs (free or associated with ribosomal subunits), monosome-associated RNAs (bound by a single ribosome), and polysome-associated RNAs (actively translated by multiple ribosomes). Because these global polysome profiles predominantly reflect the translational status of host mRNAs, which constitute the vast majority of the cellular transcriptome, potential changes in viral RNA translation cannot be inferred from global profiles alone and must be assessed by specifically quantifying viral RNAs across polysome fractions by RT-qPCR. Based on CHIKV protein expression and viral RNA level kinetics in both cell lines (**Fig 1C-E**), we selected four time points: 8 hpi, when viral protein expression is first detected; 1 dpi, corresponding to the peak of viral protein expression; and 5 and 7 dpi, when viral protein levels decline.

In U4.4, polysome profiles revealed that CHIKV infection does not induce major changes in global translation, evidenced by comparable monosome peaks (around fractions 9-12) and polysome distribution (around fractions 13-20) in infected and non-infected cells (**Fig 2A**). In contrast, C6/36 polysome profiles showed that CHIKV infection induces a transient global translational repression, evidenced by an increased monosome fraction and a reduced polysome fraction at 8 hpi (**Fig 2B**). This repression gradually diminished over time, and by 7 dpi, the translational profiles of infected and non-infected cells were similar, indicating recovery of basal translation. To quantify this effect, we calculated the polysome-to-monosome (P/M) ratio at each time point relative to non-infected controls and normalized it to 1 dpi. Values < 1 indicate a relative enrichment of monosomes over polysomes, consistent with reduced translational efficiency compared to 1 dpi, whereas values > 1 indicate the opposite. In U4.4, P/M ratios in infected cells remained comparable to those of non-infected controls throughout the time course (**Fig 2C**). In C6/36 cells, the P/M ratio indicated a global translation repression at 8 hpi. This was followed by a modest increase and a subsequent sharp increase of P/M ratios at 5 dpi before returning toward baseline levels at 7 dpi (**Fig 2D**). To investigate the molecular basis underlying the transient global translational repression observed in C6/36 but not in U4.4 cells, we examined the phosphorylation status of the eukaryotic translation initiation factor 2 subunit alpha (eIF2α) (**Fig 2E**). Under cellular stress, including viral infection, phosphorylation of eIF2α leads to attenuation of translation initiation and global repression of protein synthesis [17]. However, many viruses modulate this stress response to maintain viral protein synthesis in both vertebrate and invertebrate hosts [18–20]. Whether CHIKV induces eIF2α phosphorylation in mosquito cells, and how this response relates to viral RNA translation, remains unclear. We therefore analyzed eIF2α phosphorylation at multiple time points after infection. In U4.4 cells showed no detectable induction of eIF2α phosphorylation, consistent with the absence of global translational repression in these cells. Conversely, in C6/36 cells, a transient increase in eIF2α phosphorylation was detected between 8 hpi and 1 dpi, coinciding with the period of global host translational repression (**Fig 2E**). As infection progressed into persistence, eIF2α phosphorylation levels declined and host translation was restored.

**Fig 2.**
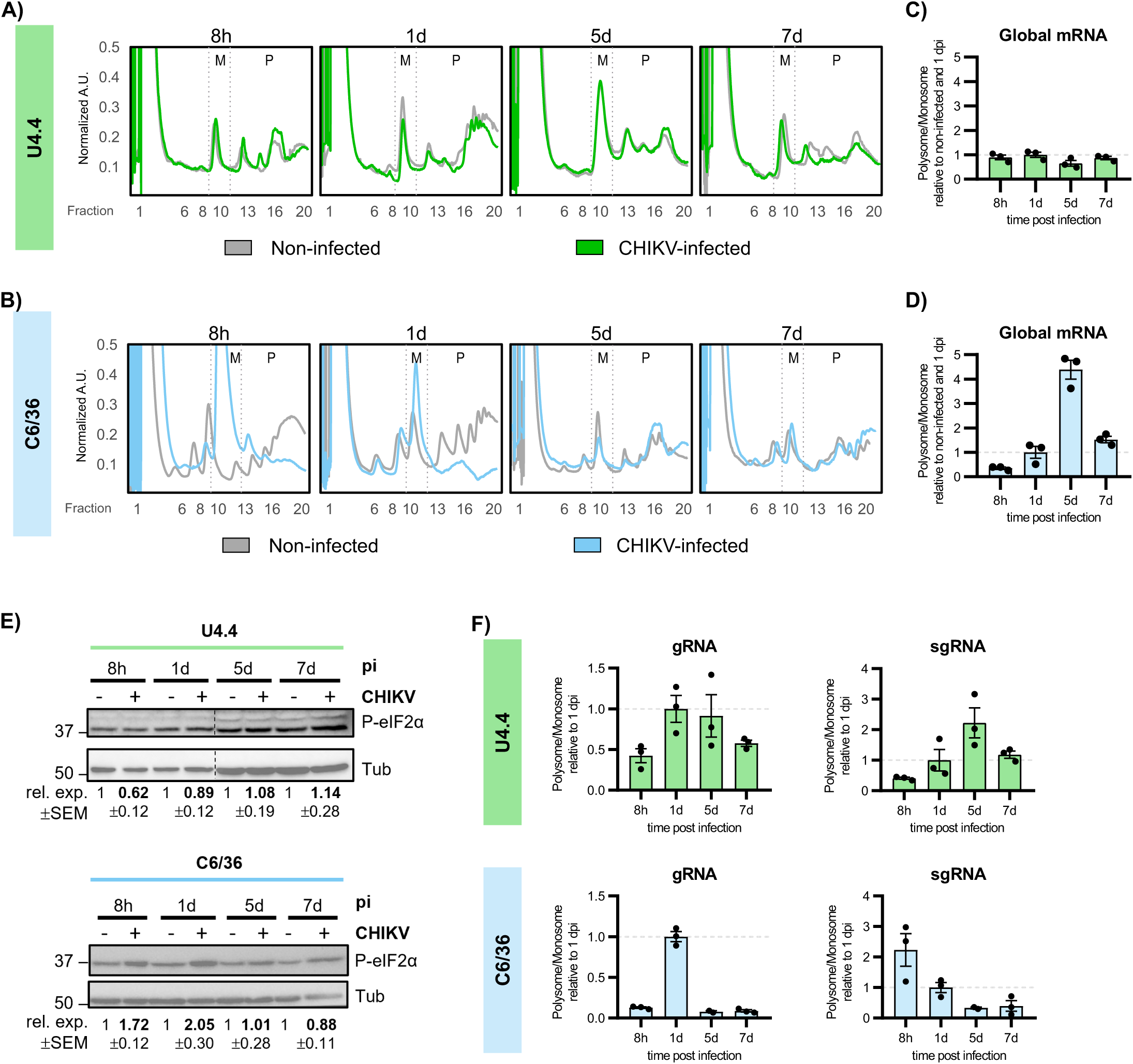
Reduced ribosome loading of viral RNAs during persistent CHIKV infection. (A-B) Representative polysome profile analyses of U4.4 cells (A) and of C6/36 cells (B) at the indicated time points after CHIKV infection. (C-D) Polysome-to-monosome (P/M) ratios reflecting global mRNA translation in U4.4 and C6/36 cells over the course of infection calculated from the area under-the-curve (AUC) of polysome profiles and normalized to non-infected controls and to 1 dpi. (E) Levels of phosphorylated eIF2α in U4.4 and C6/36 cells during CHIKV infection assessed by western blotting. Protein expression levels were normalized to tubulin and to the non-infected value within each blot. Dotted lines indicate separated lanes in the same membrane. (F) Polysome-to-monosome (P/M) ratios of viral CHIKV gRNA and sgRNA in U4.4 and C6/36 cells over the course of infection determined by RT-qPCR of RNA extracted from polysome fractions. Values are expressed relative to 1 dpi. Bars represent the mean ± SEM of 3 independent replicates.

To determine whether viral RNAs follow the same or distinct translational trends as host mRNAs, we measured the distribution of CHIKV gRNA and sgRNA across polysome fractions by RT-qPCR (**Fig 2F**). The viral P/M ratios indicated that, in both cell lines, CHIKV RNAs display translational dynamics that differ from those of the bulk host mRNAs. In U4.4 cells, the P/M ratios of gRNA and sgRNA increased up to 1 and 5 dpi, respectively, and then decreased at later time points, indicating the onset of viral RNA translation repression (**Fig 2F, green**). In C6/36 cells, gRNA followed a similar but more pronounced pattern, whereas sgRNA showed reduced association with polysomes already from 8 hpi onward (**Fig 2F, blue**). Of note, the higher P/M values for gRNA and sgRNA coincided with global host translation repression and with a transient rise in eIF2α phosphorylation between 8 hpi and 1 dpi (**Fig. 2E**). As infection progressed into the persistent phase, eIF2α phosphorylation decreased, host translation recovered, and viral RNA translation became repressed. Although viral RNA translation was repressed in both cell lines at late time points, repression appeared earlier and more evident in C6/36 cells. This is likely due to the higher viral RNA levels reached in this cell line as a result of the lack of a functional RNA interference pathway, which may increase competition between viral and host mRNAs for the cellular translation machinery, particularly after host translation resumes.

Overall, these results indicate that repression of viral RNA translation is a conserved feature of CHIKV persistence and occurs in the absence or presence of RNAi-mediated antiviral activity.

### CHIKV does not induce host transcriptional shut-off or tRNA epitranscriptome remodeling in mosquito cells

To better understand why CHIKV translation is repressed in mosquito cells, in contrast to the translational activation described during lytic infection of human cells [15], we investigated whether two key mechanisms that promote viral translation in mammalian cells are also engaged in the mosquito context. In human cells, CHIKV infection promotes viral translation through two complementary strategies. First, nuclear translocation of the viral nsP2 induces degradation of Rpb1, the catalytic subunit of RNA polymerase II. This degradation results in global host transcriptional shut-off and depletion of host mRNAs [16], thereby reducing competition between host and viral transcripts for the translation machinery. Second, CHIKV remodels the tRNA epitranscriptome, increasing the abundance of the mcm⁵ modification, thereby reprogramming codon optimality and enhancing translation of viral RNAs enriched in codons whose decoding is favored by the mcm⁵-modified tRNA modification [15].

We first examined whether the nsP2-Rpb1 axis is engaged in mosquito cells by assessing the levels of Rpb1 protein. As expected, Rpb1 levels were reduced in CHIKV-infected human HEK293T cells, whereas Rpb1 levels remained non-significantly changed in both U4.4 and C6/36 mosquito cells during CHIKV infection (**Fig 3A**). Consistently, immunofluorescence analysis showed that nsP2 remained confined to the cytoplasm in C6/36 cells (**Fig 3B**), in contrast to the established nuclear localization reported in mammalian infections [21]. nsP2 localization could not be assessed in U4.4 cells because nsP2 protein levels were below the detection limit of the immunofluorescence assay. Together, these data indicate that CHIKV does not induce nsP2-dependent host transcriptional shut-off in mosquito cells. Next, we investigated whether CHIKV modulates the mosquito translational environment through remodeling of the tRNA epitranscriptome. For this, we analyzed tRNA modification profiles at three relevant stages of infection: 8 hpi, 1 dpi, and 7 dpi. LC-MS/MS analyses revealed no significant changes in any of the monitored tRNA modifications, including mcm^5^ nor its derivative 5-methoxycarbonylmethyl-2-thiouridine (mcm^5^s^2^), in either U4.4 or C6/36 cells at any time point (**Fig 3C** and **S1 Table**).

**Fig 3.**
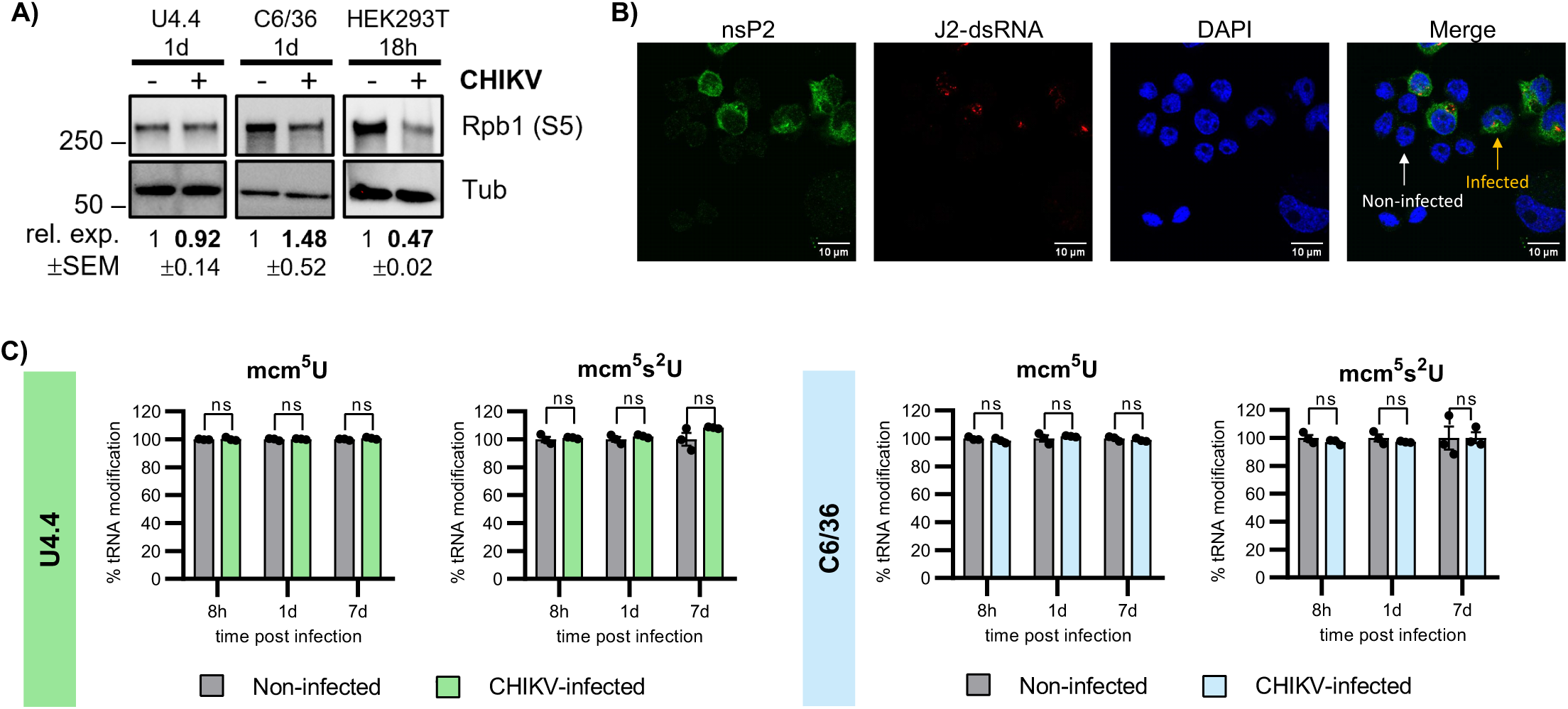
CHIKV does not induce Rpb1 degradation or tRNA epitranscriptome remodeling in mosquito cells. (A) Western blot analysis of Rpb1 protein levels in U4.4, C6/36, and HEK293T cells non-infected and infected with CHIKV. Protein expression levels were normalized to tubulin and to the non-infected value within each blot. (B) Immunofluorescence analyses of nsP2 subcellular localization in CHIKV-infected C6/36 cells 1 dpi. Cells were labeled with anti-CHIKV nsP2 antibody (green), J2-dsRNA (red), and nuclei were stained with DAPI (blue). Scale bars correspond to 10 μm. (C) LC–MS/MS quantification of mcm⁵U and mcm⁵s²U tRNA modifications in CHIKV-infected U4.4 and C6/36 cells. Values are expressed relative to those in non-infected. Bars represent the mean ± SEM of 3 independent replicates.

Collectively, these results indicate that two key CHIKV strategies that promote viral translation during lytic infection of human cells, host transcriptional shut-off and tRNA epitranscriptome remodeling, are not engaged in mosquito cells.

### Impaired viral protein production is a conserved feature of arbovirus infection in mosquito cells

To determine whether the viral RNA translational repression observed for CHIKV represents a conserved mechanism among arboviruses in mosquito cells, we extended our analysis to ZIKV. ZIKV is a (+)ssRNA flavivirus with a ∼10.7 kb genome bearing a 5’ cap but lacking a 3’ poly(A) tail. The ZIKV genome contains a single ORF that is translated into a polyprotein, which is subsequently processed into individual structural and non-structural (NS) proteins (**Fig 4A**). U4.4 cells were infected with ZIKV at an MOI of 1, and infection kinetics were monitored over a 13 day-period. Viral titers (**Fig 4B**), viral RNA levels (**Fig 4C**) and NS3 protein expression (**Fig 4D**) were quantified at 1, 4, 7, 10, and 13 dpi. Viral titers increased until 7 dpi and subsequently slightly declined. In contrast, viral RNA levels peaked at 4 dpi, showed a modest decrease up to 10 dpi, and increased again at later time points.

**Fig 4.**
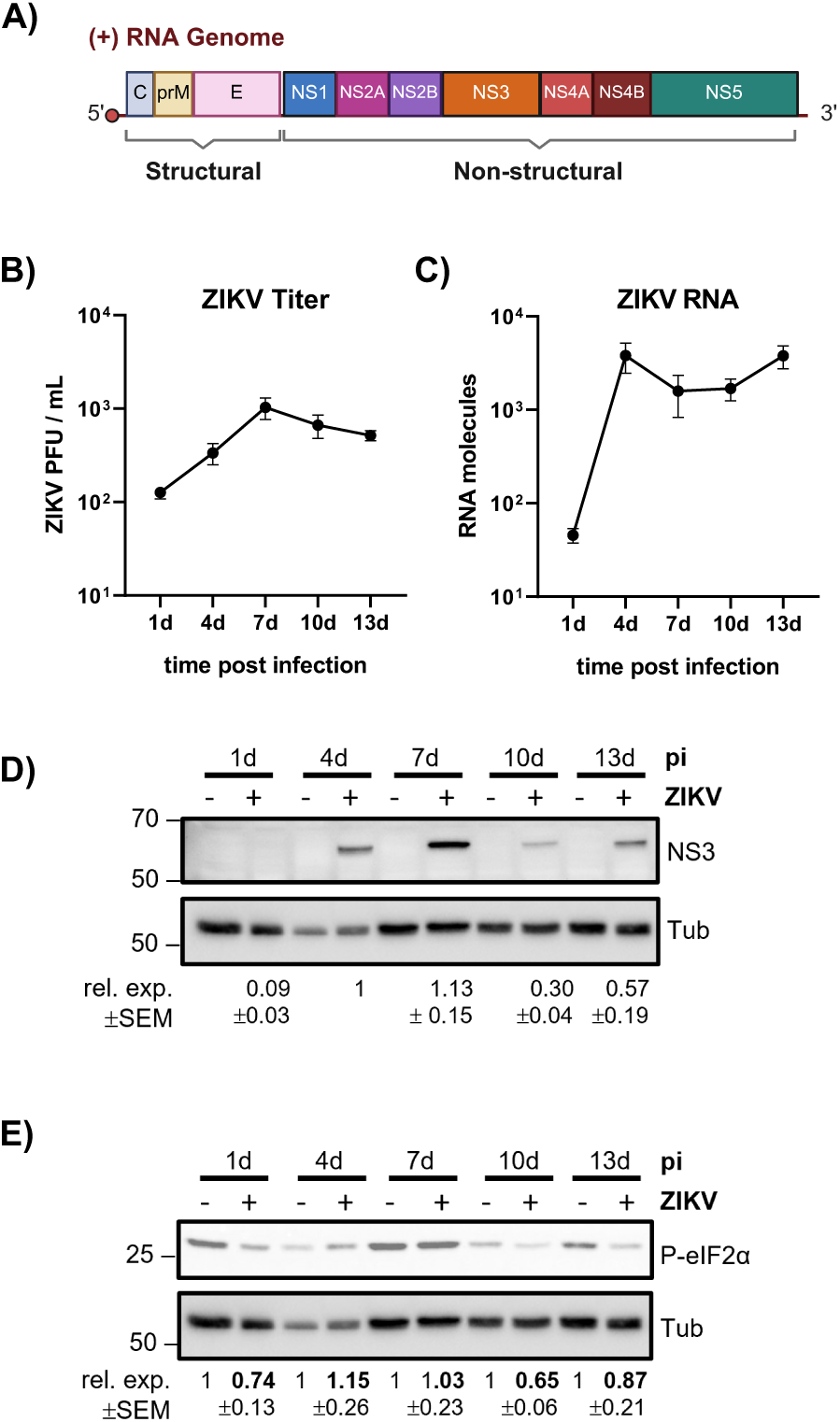
Kinetic analysis of ZIKV infection in U4.4 mosquito cells. (A) Scheme of ZIKV RNA genome and its protein synthesis strategy. (B) ZIKV titers over time, determined by plaque assay. (C) Infection kinetics of ZIKV RNA levels determined by qPCR. Data points represent the mean ± SEM of 3 independent replicates. Viral RNA levels were quantified using a standard curve of an *in vitro* generated ZIKV RNA. (D) NS3 protein expression levels assessed by western blotting. Protein expression levels correspond to the intensity quantification of each band normalized to tubulin levels and to the 4 dpi value within each blot. (E) Levels of phosphorylated eIF2α protein levels during U4.4 ZIKV-infection, assessed by western blotting. Protein expression levels correspond to the intensity quantification of each band normalized to tubulin and to the non-infected value within each blot. Infection was performed at an MOI of 1 and samples were collected at the indicated time post infection.

Importantly, ZIKV NS3 protein peaked at 7 dpi and then declined, despite sustained or increasing viral RNA levels. Thus, ZIKV infection in mosquito cells displays a discordance between viral RNA accumulation and viral protein production, similar to that observed for CHIKV. To exclude the possibility that this reduction in viral protein production reflects a general host stress response rather than a virus-specific translation regulatory mechanism, we examined the phosphorylation status of eIF2α in ZIKV-infected cells. No increase in eIF2α phosphorylation was detected compared with mock-infected controls (**Fig 4E**).

Together, these data show that ZIKV, like CHIKV, exhibits reduced viral protein production during persistence in mosquito cells in the absence of detectable activation of the integrated stress response. This supports the notion that regulated limitation of viral protein synthesis is a common feature of arbovirus persistence in mosquito cells

## Discussion

Arboviruses establish long-term, non-cytopathic infections in mosquito vectors, thereby helping to ensure efficient transmission, yet the molecular mechanisms that enable this balance remain poorly understood. Here, using CHIKV infection of *Aedes albopictus* U4.4 and C6/36 cells as a model, we identify translational control as a characteristic feature of viral persistence in mosquito cells. Persistent infection is marked by an uncoupling between viral RNA accumulation and viral protein production, driven by a virus-specific repression of viral RNA translation. This translational restriction occurs independently of the RNAi antiviral pathway and establishes a finely tuned virus-host translational equilibrium that supports sustained viral genome maintenance while preserving host cell viability. This behavior contrasts sharply with the highly efficient viral protein synthesis observed during lytic infection of human cells, in which CHIKV actively remodels the host translational environment to favor viral gene expression [15]. Collectively, our data indicate that, rather than maximizing viral output, CHIKV adopts a self-limiting translational strategy in mosquito cells that is compatible with long-term infection.

Our results further identify two major host-specific determinants that shape viral translational efficiency in human versus mosquito cells: codon optimality and competition for translational resources (**Fig 3**). CHIKV genomes are intrinsically enriched in codons that are suboptimal in both hosts. In human cells, however, CHIKV infection induces remodeling of the tRNA epitranscriptome, thereby reprogramming codon optimality in a manner that favors viral RNA translation [15]. In contrast, mosquito cells do not remodel their tRNA modification landscape during infection and CHIKV codon usage therefore remains suboptimal throughout the viral cycle. In parallel, whereas CHIKV infection in human cells causes a marked depletion of host mRNAs through nuclear relocalization of the viral protein nsP2, it remains cytoplasmic in mosquito cells. The combination of continued suboptimal codon decoding and sustained competition with a stable pool of host mRNAs are therefore expected to constrain viral protein synthesis in mosquito cells and represent a major mechanistic difference between lytic infection in humans and persistent infection in mosquitoes (**Fig 5**).

**Fig 5.**
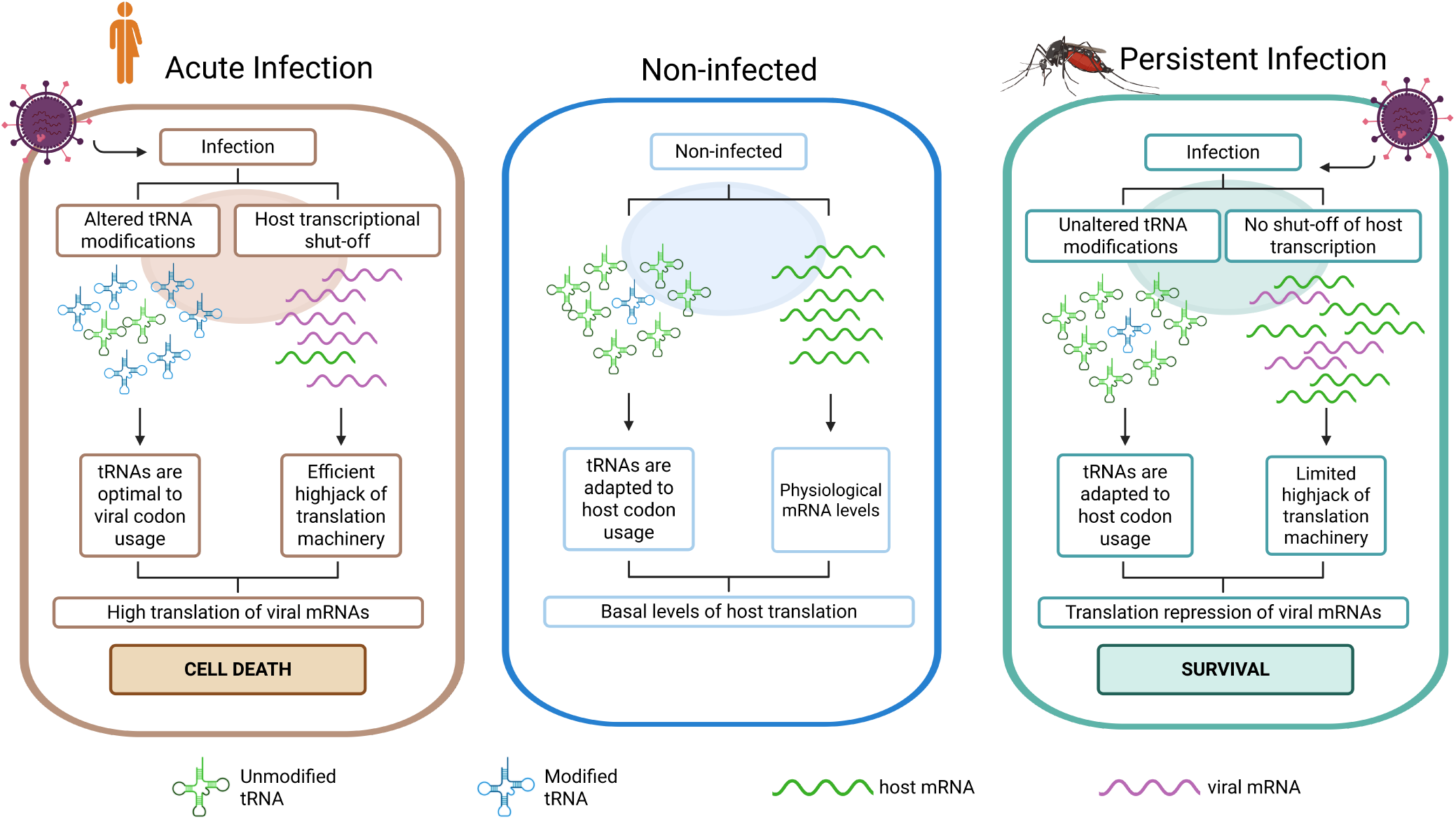
Model of arbovirus infection in humans and mosquitoes. In non-infected human and mosquito cells, tRNAs are adapted to host codon usage and mRNA levels are maintained to meet cellular needs. During acute CHIKV infection in human cells, remodeling of the tRNA epitranscriptome favors viral codon usage and host mRNA levels are strongly reduced, resulting in highly efficient viral translation and lytic infection. In contrast, during persistent infection in mosquito cells, neither the tRNA epitranscriptome nor host mRNA levels are altered, viral codons remain suboptimal for translation, and viral protein synthesis is limited despite sustained viral RNA levels, allowing long-term survival of infected cells.

In RNAi-deficient C6/36 cells, but not in RNAi-competent U4.4 cells, we observed a transient phosphorylation of eIF2α that coincided with host translation repression and viral translation activation (**Fig 2E**). During this early phase, viral RNAs, particularly sgRNAs, remain efficiently translated, indicating that CHIKV can maintain viral protein synthesis in the context of eIF2α phosphorylation-mediated translational repression. This property is consistent with observations in other alphaviruses, such as Sindbis and Semliki Forest viruses, whose subgenomic RNAs harbor structural features that support translation initiation under conditions of eIF2α phosphorylation [22]. We speculate that the ability of CHIKV to sustain translation under eIF2α phosphorylation may provide an advantage during acute infection in human cells, where viral replication is rapid and activation of the integrated stress response is more likely. In contrast, in mosquito cells, where viral replication is limited by RNAi and eIF2α phosphorylation is not observed, this stress-resilient mode of viral translation would be less critical. As infection progresses toward persistence in C6/36 cells, recovery of host translation and decline of eIF2α phosphorylation coincide with progressive repression of viral RNA translation, further reinforcing a transition to a competitive environment for translation resources.

Importantly, this translation-restricted phenotype is not limited to CHIKV. ZIKV infection of mosquito cells likewise displays a discordance between viral RNA accumulation and viral protein production, in the absence of detectable activation of eIF2α phosphorylation. In vertebrate cells, ZIKV promotes host shut-off and remodels the tRNA epitranscriptome [23–25]. This enhances viral translation, indicating that, as for CHIKV, host-directed mechanisms that optimize viral gene expression in vertebrate hosts would not be engaged in mosquito cells. The conservation of viral translational repression across distinct arboviruses therefore suggests that controlled limitation of viral protein synthesis may represent a general outcome of long-term co-evolution between arboviruses and their mosquito vectors.

In summary, our findings identify regulation at the level of translation as a defining feature of arbovirus persistence in mosquito vectors and underscore translational control as a major point of divergence between arbovirus infection in vertebrate and invertebrate hosts.

## Materials and Methods

### Cell Lines and Viral stocks

C6/36 (kindly provided by Andres Merits) and U4.4 (kindly provided by Emilie Pondeville) cells were grown at 28°C and cultivated in Leibovitz 15 medium (L15; ThermoFisher, 11415064) supplemented with 10% Tryptose Phosphate Broth (TPB; BD, 260300) and 10% Fetal Bovine Serum (FBS; Merck, F7524-500ML). HEK293T (ATCC, CRL-11268) cells were grown at 37°C and 5% CO_2_ and cultivated in Dulbecco’s modified Eagle’s medium (DMEM; ThermoFisher Scientific, 41966029) supplemented with 10% FBS and 1% non-essential amino acids (ThermoFisher Scientific,11140035).

Stock of CHIKV LR2006-OPY1 (GenBank: DQ443544, kindly provided by Andres Merits) was generated as in [26]. For production of ZIKV stock (Asian genotype, isolate BeH819016; GenBank: KU365778.1, kindly provided by Andres Merits), 8x10^6^ Vero cells were resuspended in 800 μl of PBS and electroporated with 7 μg of *in vitro* ZIKV generated as in [26] in an electroporation cuvette (0.4 cm gap). Cells were then pulsed three times at 450 V, 25 μF with a Gene Pulser Xcell system (Bio-Rad) and directly resuspended in DMEM supplemented with 10% FBS and 1% non-essential amino acids. ZIKV-containing supernatant was collected every 72 h post-electroporation until cytopathic effect was observed. Viral stock was filtered with a 0.45 μm filter and mixed with all previous aliquots before titration. Virus titers were determined by plaque assay using HEK293T cells as described below.

### Viral infections

All CHIKV infections were carried out with an MOI of 4 and incubated for 1 h. ZIKV infection was carried out with an MOI of 1 and incubated for 4 h. Samples were harvested at the indicated time points after infection.

For all C6/36 infections, plates were previously coated with poly-L-lysine (Merck, P1274-100MG). 6-well plates were seeded with 5x10^5^ C6/36 cells/well or 3x10^5^ U4.4 cells/well. On the next day, cells were washed once with PBS (Cytiva, SH30028.02) and then infected with viral stock. After incubation, the virus was removed, and 2 mL of pre-warmed media was added. At the time of harvest, supernatant was taken and stored at -80°C until titration; cells were washed once with PBS and then lysed with either 250 µl ice-cold lysis buffer (10 mM Tris-HCl pH 7.4, 10 mM MgCl_2_, 100 mM NaCl, 1% Triton, 2 mM DTT) for RNA extraction or 200 μl of lammeli buffer for protein analyses. For polysome profiling, we seeded 1x10^7^ C6/36 cells/T-150 and 3x10^6^ U4.4 cells/T-150. At the time of harvest, cells were previously incubated with non-supplemented media with 100 μg/mL cycloheximide (CHX; Merck, C7698), then washed once with PBS + 100 μg/mL CHX and finally lysed with 700 μl ice-cold lysis buffer + 0.25% deoxycholate and 100 μg/ml CHX. Cells were scraped and transferred to a new tube and snap-frozen with liquid nitrogen. For time points longer than 1 dpi, C6/36, cells were jumped out in fresh media at the time of harvest, and U4.4 were trypsinized for 5 min (ThermoFisher Scientific, 25200056). Then, 1:5^th^ was re-plated in new plates.

Cell viability was tested by following both cell growth and accumulation of intracellular ATP levels using CellTiter-Glo (Promega, G7570).

### RNA extraction and RT-qPCR analyses

For further processing, all lysates were centrifuged at 4°C for 5 min at 12.000 xg and transferred to new tubes for subsequent analysis. Total RNA was isolated following proteinase K treatment (New England Biolabs, P8107S), phenol-chloropropane extraction, and ethanol precipitation. The pellet was resuspended in RNAse-free water and consecutively treated with 0.1 µl Turbo DNase/μg RNA (ThermoScientific, AM1907) for 30 min at 37°C. 18 nanograms (ng) of total RNA was analyzed by TaqMan RT-qPCR using qScript XLT One-Step RT-qPCR ToughMix (Quanta BioSciences, 733-2232). The following primers and probes were used: CHIKV gRNA Fwd (5’-aaccccgttcatgtacaatgc-3’), Rev (5’-gtacctgctcatctgcccaatt-3’) and probe (5’-6Fam-cgggtgcctacccctcatactcgac-TAMRA-3’); CHIKV sgRNA Fwd (5’-aagctccgcgtcctttaccaag-3’), Rev (5’-ccaaattgtcctggtcttcct-3’) and probe (5’-6Fam-ccaatgtcttcagcctggacaccttt-TAMRA-3’); ZIKV Fwd (5’-gtgcaacacgacgtcaacttg-3’), Rev (5’-ggtttgcgaccgcgttt-3’) and probe (5’-6Fam-agagctgtgacgctcccctccca-TAMRA-3’). For the RT-qPCR of the CHIKV (-)gRNA we adapted the protocol described in [27] considering the viral mutations present in the strain used in this work. The following primers and probes were used: CHIKV Fwd T tag T (for RT) (5’-ggcagtatcgtgaattcgatgcgacacggagacgccaacatt-3’), CHIKV Rev T (5’-aataaatcataagtctgctctctgtctacatga-3’), CHIKV (-)gRNA probe (5’-6Fam-tgcttacacacagacgt-TAMRA-3’), Tag T (5’-aataaatcataaggcagtatcgtgaattcgatgc-3’).

For the standard curve of (+)RNA *in vitro* CHIKV and ZIKV were generated as in [21] and were serially diluted 1:10 to generate a standard curve for absolute quantification of viral RNA molecules. To generate the standard curve of the (-)gRNA, the same protocol was performed but using the following primers: nsP1 Fwd (5’-atcatggatcctgtgtacgtggac-3’) and nsP1 Rev T7 (5’-taatacgactcactatagtgcgcccgtctggtcctcaa-3’).

### Protein analyses

Lammeli lyzed samples were boiled for 5 min at 95°C and subsequently loaded onto a 6% or 10% polyacrylamide-SDS gel. Following electrophoresis, proteins were transferred to a membrane using TurboBlot (Biorad) for 7-11 min at 25V. Then the membranes were blocked with 5% non fat dry milk in TBST for 1 h at room temperature and then incubated overnight at 4°C with the primary antibodies. Antibodies used in western blotting were: CHIKV nsP1 (1:1500), CHIKV Capsid (1:1500), CHIKV nsP2 (1:5000) and ZIKV NS3 (1:2000), kindly provided by Andres Merits; Tubulin (1:1000; Merck, T6199-100UL), Rpb1 (1:1000; Active Motif, 39497) and P-eIF2α (1:1000; Cell Signaling, 9721S). Next day they were washed 3x for 5 min with TBST and incubated at room temperature for 1 h with secondary antibodies (GE Healthcare, NA934V or NA931V) diluted in 5% milk in TBST. Subsequently, the membranes were washed again 3x for 5 min with TBST and exposed to ECL solution (Cytiva, GERPN2134D2 or ThermoFisher, 10005943). Band detection was done using the BioRad ChemiDoc MP Imaging System. Total band intensity volumes of proteins of interest were normalized to the housekeeping gene of the same sample, and the mean of 3 biological replicates was calculated. Viral protein levels were further normalized to the highest intensity time point (1 dpi for CHIKV and 4 dpi for ZIKV). P-eIF2α in infected samples was normalized to the mock levels of the protein at each corresponding time point.

### Viral Titer

Supernatants were titrated using standard plaque assay. Briefly, supernatants for each time point were diluted 1:10 in non-supplemented DMEM media. HEK293T cells were seeded at 3x10^5^ cells/well in 12-well plates and incubated with the diluted supernatants for 1 h at 37°C and shaken every 10 min. After incubation, the virus was changed to pre-warmed DMEM + 2% FBS + 1% carboxymethyl cellulose (CMC; Merck, 21902) in a 1:1 ratio and put back into a 37°C incubator for 3 days (CHIKV) or 7 days (ZIKV). Then, cells were fixed and stained with crystal violet for 30 min under UV light. Finally, cells were rinsed with warm tap water, and plaques were counted.

### Immunofluorescence

For immunofluorescence analysis, we seeded 2x10^5^ cells/well for C6/36 and 8x10^4^ cells/well for U4.4 into 24-well plates containing sterile glass coverslips coated with poly-L-lysine. Cells were infected with CHIKV as explained before, and at different time points post infection, cells were processed for confocal study. Cells were washed with PBS and fixed with 4% paraformaldehyde (Sigma-Aldrich, P6148) for 15 min. Then, they were permeabilized with PBST (PBS + 0.5% Triton X-100; Sigma-Aldrich, X-100) for 20 min in ice. After a wash with PBS, cells were blocked in blocking buffer (PBS + 10% FBS) for 30 min at room temperature, washed, and then incubated for 1 h with the different antibodies. Using blocking buffer, antibodies were diluted at: 1:10000 for anti-Capsid (rabbit) and 1:1000 for anti-nsP2 (rabbit), kindly provided by Andres Merits, and 1:2500 for anti-J2 (mouse) and anti-double-stranded RNA (clone J2) (Nordic-MUbio, 10010200). Then, cells were washed twice with PBS and incubated with the secondary antibody anti-rabbit Alexa FluorTM 488 goat IgG and anti-mouse Alexa FluorTM 568 goat IgG (ThermoFisher, A-11004 and -11008) diluted 1:1000 in blocking buffer for 45 min covered from the light. Cells were washed and immersed in DAPI staining (1 µg/ml, Sigma-Aldrich) for 5 min, covered from the light. Lastly, slides were mounted using pre-warmed Mowiol (Sigma-Aldrich, 32459). Cells were imaged using a Leica SP5 inverted confocal laser scanning microscope and processed with Fiji (ImageJ). For percentage calculations, 4 randomly selected fields per slide were acquired from at least two biological replicates for each infection time point.

### Polysome Profiling

Sucrose (Sigma-Aldrich, 84097) solutions at 10% and 50% were prepared dissolved in polysome buffer 2.0 (20mM Tris-HCL pH 7.5, 10mM MgCl2, 100 mM NH4Cl), and supplemented with CHX 100ug/ml and DTT 2mM. 6 mL of the 10% sucrose gradient was added to SW41Ti tubes (Beckman, 344059) and then 6 mL of 50% sucrose was underlaid using a needle. To get a 15-45% linear gradient, we used the Gradient Master. Gradients were stored at 4°C overnight. Lysates were thawed on ice and centrifuged for 5 min at 12.000 xg. 400-700 µL of lysate coming from the same number of cells, were loaded onto the linear sucrose gradients, and centrifuged for 3 h at 35,000 rpm at 4°C in a Beckman SW41 rotor. BCA quantification was performed to normalize the amount of sample loaded into the gradient. Gradients were immediately fractionated into 20 fractions using a Biocomp Fractionator. Fractions of 500 µl were collected in 1.5 mL RNAse-free tubes with 55 µL of 10% SDS.

Fractions were extracted using phenol-chloroform and precipitated with ethanol. RNA of fractions corresponding to 40S, 60S, 80S, and polysome peaks were pooled together when dissolving with RNAse free water. Samples were DNase-treated before analyzing by qPCR. Viral RNA was assessed as described above using TaqMan qPCR.

### Quantification of tRNA modifications by LC-MS/MS

3-10 μg of total RNA per sample was run in a 15% TBE-UREA gel (Novex, ThermoFisher Scientific, EC6885BOX) for 60 min at 180 V Gels were stained with 1:10000 SybrGold (ThermoFisher Scientific, S11494) and the tRNA bands were selected. tRNA was then extracted using the ZR small RNA PAGE Recovery kit (Zymo Research, R1070) following the manufacturer’s instructions. The same amount of extracted tRNA was digested with 1 μl of the Nucleoside Digestion Mix (New England BioLabs, M0649S) and the mixture was incubated at 37°C for 1 h. Samples (15 ng) were analyzed using an Orbitrap Eclipse Tribrid mass spectrometer (ThermoFisher Scientific, San Jose, CA, USA) coupled to an EASY-nLC 1200 (ThermoFisher Scientific (Proxeon), Odense, Denmark). Ribonucleosides were loaded directly onto the analytical column and were separated by reversed-phase chromatography using a 50 cm column with an inner diameter of 75 μm, packed with 2 μm C18 particles (ThermoFisher Scientific, ES903). Chromatographic gradients started at 93% buffer A and 3% buffer B with a flow rate of 250 nL/min for 5 min and gradually increased to 30% buffer B and 70% buffer A in 20 min. After each analysis, the column was washed for 10 min with 0% buffer A and 100% buffer B. Buffer A: 0.1% formic acid in water. Buffer B: 0.1% formic acid in 80% acetonitrile. The mass spectrometer was operated in positive ionization mode with nanospray voltage set at 2.4kV and source temperature at 305°C. For Parallel Reaction Monitoring (PRM) method the quadrupole isolation window was set to 1.4m/z, and MSMS scans were collected over a mass range of m/z 50-300, with detection in the Orbitrap at a resolution of 60,000. MSMS fragmentation of defined masses (**S1 Table**) was performed using HCD at NCE 20 (except stated differently) [28], the auto gain control (AGC) was set to “Standard” and a maximum injection time of 118 ms was used. In each PRM cycle, one full MS scan at a resolution of 120,000 was acquired over a mass range of m/z 220-700 with detection in the Orbitrap mass analyzer. Auto gain control (AGC) was set to 10^5^ and the maximum injection time was set to 50 ms. Serial dilutions were prepared using commercial pure ribonucleosides (0.005-150 pg, Carbosynth, Toronto Research Chemicals) in order to establish the linear range of quantification and the limit of detection of each compound. A mix of commercial ribonucleosides was injected before and after each batch of samples to assess instrument stability and to be used as an external standard to calibrate the retention time of each ribonucleoside.

Acquired data were analyzed with the Skyline-daily software (v24.1.1.284) and extracted precursor areas of the ribonucleosides were used for quantification.

## Data availability

The raw proteomics data have been deposited to the EMBL-EBI MetaboLights database with the identifier REQ20250616211235.

## Acknowledgements

This work has received funding by “la Caixa” Foundation (LCF/PR/HR23/52430003), the Spanish Ministry of Science and Innovation (PID2022-136939OBI00/MICIN/AEI/10.13039/501100011033 and “ERDF a way of making Europe”), the 2021 SGR 00176 grant from the Departament de Recerca i Universitats de la Generalitat de Catalunya, and by an institutional “Unidad de Excelencia María de Maeztu,” CEX2024-001431-M, funded by the MICIU/AEI/10.13039/501100011033. MP-T was a recipient of an FPI fellowship from the Spanish Ministry of Science and Innovation (PRE2020-093049). We are grateful to the UPF and CRG Core Technologies Programmes for their support and assistance in this work. We thank R. Barrios for insightful discussions and E. Muscolino and A. Meyerhans for critically reading the manuscript and for insightful discussions. Figures 1A, 4A and 5 were created using BioRender for which we own a full license.

## Supporting Figures/Tables Captions

**S1 Fig.**
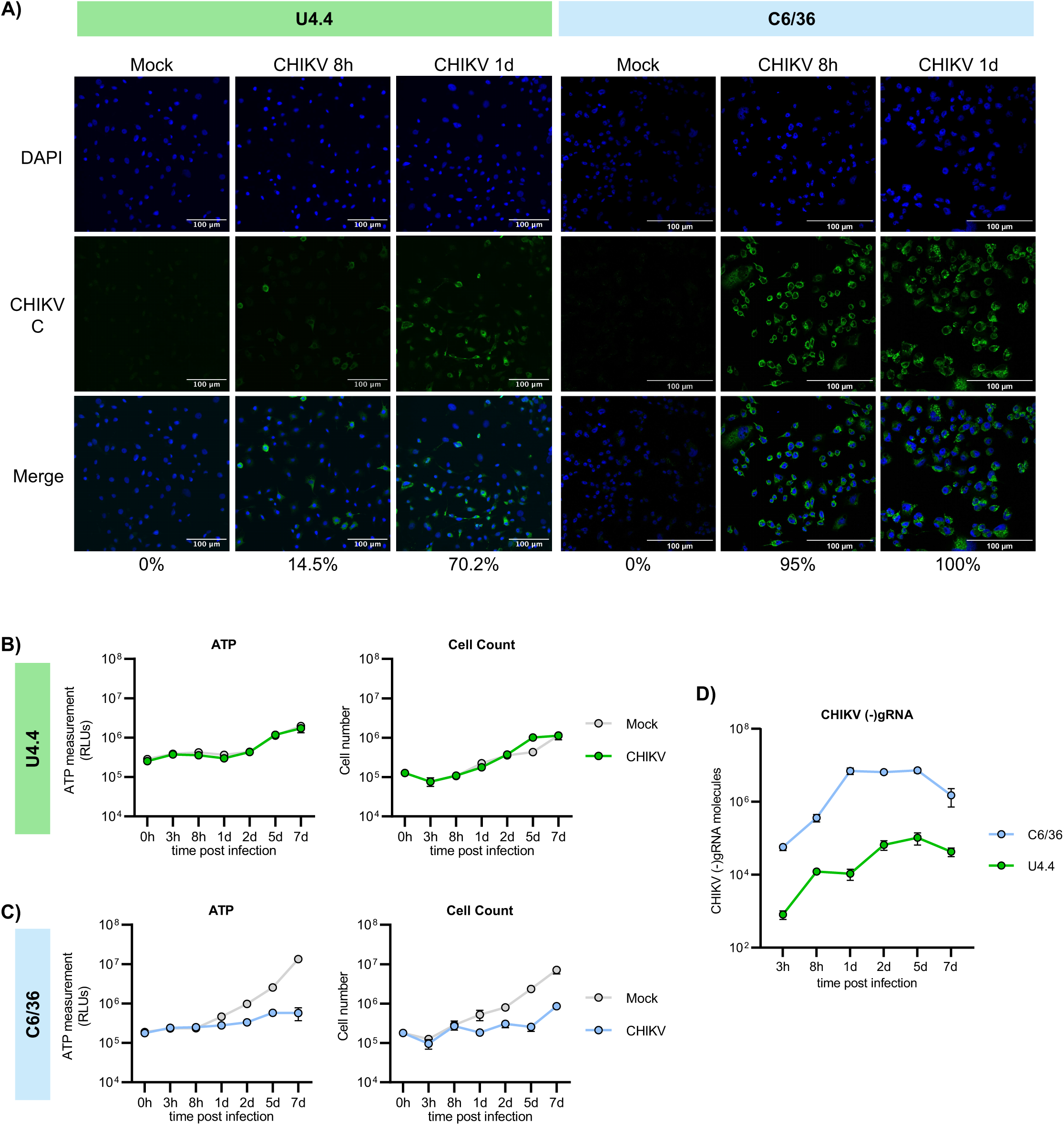
Infection dynamics, cell proliferation and metabolic activity during CHIKV infection in U4.4 and C6/36 cells. (A) Immunofluorescence analysis of CHIKV infection in U4.4 and C6/36 cells at the indicated time points. Cells were labeled with anti-CHIKV capsid antibody (green), and nuclei were stained with DAPI (blue). Scale bars correspond to 100 μm. Percentage indicates the proportion of cells positive for the capsid CHIKV-protein. (B-C) Cell counts and cellular ATP levels in CHIKV-infected and mock-infected U4.4 and C6/36 cells over time. (D) Kinetics of CHIKV negative-strand RNA ((-)gRNA) levels in U4.4 and C6/36 cells, quantified by RT-qPCR. Blue and green colors correspond to C6/36 and U4.4 cells, respectively. All infections were performed at a MOI of 4 and samples were collected at the indicated times post infection. Data points represent the mean ± SEM of 3 independent replicates. Negative-strand viral RNA levels were quantified using a standard curve generated from an *in vitro* generated CHIKV (-)gRNA.

**S1 Table. List of tRNA modifications.** List of all modified ribonucleosides identified in mass spectrometry analyses of tRNA samples with defined masses of its MS/MS fragmentation.

